# Linking protein residues in literature and structure

**DOI:** 10.1101/2025.10.17.683004

**Authors:** Melanie Vollmar, Simon Westrip, Sreenath Nair, Balakumaran Balasubramaniyan, Sameer Velankar, Louise Jones, Peter Strickland

## Abstract

Protein structures are crucial in understanding function, mechanism and disease-causing variants of proteins within any living cell. A number of experimental techniques are employed by researchers to determine said structure. Through structure inspection in molecular viewers combined with supporting biochemical and biophysical experiments, scientists are able to identify a protein’s function, reaction mechanism and effects caused by sequence variation. These detailed findings supported by experimental results are documented and described in detail in scientific literature and by open sourcing the accompanying data. By writing a detailed report about the findings and providing evidence in additional files and complementary data formats it has become increasingly difficult for a reader, in particular a non-expert, to access the correct additional information and assess the validity of the drawn conclusion based on experimental results. It often requires a reader to resort to a number of different software packages to access the different data types. Here, we present a first-of-its-kind implementation of an artificial intelligence and text mining supported software tool that allows linking of text mentions of specific protein residues to their corresponding counterpart in the respective protein structure. An identified residue is highlighted in the publication text and upon interaction with the annotation, a molecule viewer displays the associated protein structures in the publication which contain said residue. The viewer is complemented by a display table that contains protein structure quality metrics for each occurrence of a residue. As such a reader can now explore a residue of interest they are currently assessing in a publication within its respective protein structure supported by its experimental evidence in a single view and application.

## 1. Introduction

Writing scientific publications is a well established practice in research to document and disseminate novel findings and provide background and details of the research undertaken. Written documentation ensures findings are preserved for future reference and additional investigation. Documenting experimental details also enables other researchers to cross-examine and even reproduce experiments.

In structural biology, scientific publications focus on the mechanistic understanding of macromolecular function and describe residue-level, atomic details identified through analysis of a protein structure. The experimental techniques employed to determine a protein structure, such as cryoEM, NMR, or X-ray diffraction, can vary, and the findings suffer from different types of uncertainties and errors. However, the analysis and interpretation of protein structures follow a similar process:

- Setting the scene: describe a structure’s overall architecture, secondary structure elements, domain boundaries, motifs and other details from existing literature.
- Description of the experiment quality: analyse a structure in isolation and evaluate its quality based on established metrics.
- Identify novel findings: compare a structure to related models from other organisms or in a different state to gain mechanistic and functional insights.

To facilitate comparative analysis and gain novel insights from a protein structure, a researcher must familiarize themselves with relevant literature and also consider deriving conclusions that take into account the experimental setup, data quality, and results. wwPDB (Berman *et al*., 2003) provides a validation report at the time of structure deposition in the core structure archives PDB (wwPDB consortium, 2019) or EMDB (wwPDB consortium, 2024). The validation reports are tailored specifically for each experimental technique used in structure determination, whether X-ray crystallography, cryo-electron microscopy or nuclear magnetic resonance spectroscopy and were designed by community experts as part of the validation task forces (Read *et al*., 2011; Henderson *et al*., 2012; Montelione *et al*., 2013; Baskaran *et al*., 2024). These reports provide valuable information on the overall quality of experimental data and the structure model, as well as detailed quality information on a per-residue basis. Expert users can view these validation data using specialized 3D molecule viewers. Knowledge and findings about the structure are accessed through scientific publications, which can be viewed via web-based browsers or require a PDF viewer to display the manuscript. Providing easy access to experimental evidence at a time when publications are accessed on a journal website can facilitate interpretation of biological insights presented in the publication for both structural and non-structural biologists, and support the use of structure data to drive life science research. It is especially important to ensure that mentions of specific residues with functional roles are supported by experimental evidence, thereby allowing for confident planning of further experimental studies using this information.

In the past, attempts were made to identify functionally important residue mentions in literature using rule-based text mining approaches, such as in *pyresid* (Firth *et al*., 2019) or mutation grounder (Laurila *et al*., 2010; Klein *et al*., 2012). These systems all adhere to a similar sequential execution of entity extraction, generation of candidate lists of named entities, followed by filtering and selecting best fitting candidates based on expert-defined criteria. *pyresid* is an application co-developed between EMBL-EBI and the Science, Technology and Facilities Council (STFC) aimed at identifying mentions of amino acids in publications and linking them to their respective protein. Mutation grounder was developed to identify amino acid mutations in publications and link them to reference sequence positions and accessions in UniProt (The UniProt Consortium, 2025). Graph-association rules were used by Ravikumar *et al*. (2012) which allowed for contextualising residue-protein pairs and encoding their relationship across longer distances beyond sentence boundaries.

Although these systems represent initial attempts at curating unstructured information from scientific literature into structured knowledge, they were all limited by the fine-grainedness of the decision rules that could be crafted. None of these systems was context-aware and was prone to language-ambiguity-related misidentification of relevant details.

The rapid developments in the field of natural language processing (NLP) and text mining, in particular applications for transformer and (large) language models, have made it possible to process large amounts of unstructured scientific literature. The extracted knowledge can be structured in open-access databases and made available to a wide range of scientific communities. Using structured knowledge as a linker between scientific publication and the display of a protein structure, written statements and conclusions about a specific residue in a protein structure can be placed in the experimental context. A collaboration between IUCr and PDBe has resulted in such an application.

We present here the details of a novel application that links residue mentions in publications in the IUCr journals, Acta Cryst. D, Acta Cryst. F and IUCrJ, in particular, to their respective residues in PDB structures, alongside data quality statistics from experimental results. The application uses a fine tuned transformer model described in Vollmar *et al*. (2024) to perform named entity recognition to identify any mentions of a protein residue in the format amino acid name and sequence position, as well as point mutations. Amino acids can be identified in both three- and single-letter notation. Downstream pipelines utilize SIFTS-mapping (Structure Integration with Function, Taxonomy, and Sequence; Dana *et al*., 2019) to map UniProt reference sequences and validation files provided by PDBe (Armstrong *et al*., 2020) for each structure linked to a publication, to pair the residue identified in the literature with its equivalent in the structure and its corresponding reference sequence in UniProt. This application enables readers to view a protein structure and access validation information for specific residues while reading the scientific publication. Furthermore, the harvested residue-structure-publication links are returned to PDBe and are included in the resource, where the details can be accessed through application programming interfaces (APIs).

In future implementations, we envisage placing the tool within the review pipeline to make it available to reviewers and editors at the point of new publication submissions. It is hoped that an in-line link between text mentions, structure display and experimental statistics will support reviewers and accelerate the review process. Another potential future extension could be to not only focus on individual residues and point mutations but also provide grounding and linking of additional text mentions, for example, a protein’s name, protein family, its functional state and cellular location. Our underlying predictive model already identifies additional terms. However, existing reference sources, such as ontologies and controlled vocabularies, often cover only a portion of the required terms, and initial work will be required to expand their term coverage to serve as a comprehensive reference.

## 2. Materials and Methods

### 2.1. Implementation

The transformer model, v2.1, underpinning the named entity recognition (NER) step has been described previously (Vollmar *et al*., 2024) and can be freely accessed and downloaded from https://huggingface.co/PDBEurope. A Pubmed-BERT (Gu *et al*., 2021) model was fine-tuned to carry out NER for structure specific terminology. 20 different entity types are recognized by the model with differing accuracies. The two entity types used for developing the linking pipeline are ‘residue_name_number’ with precision 0.95, recall 0.96 and F1 score 0.96, and ‘mutant’ with precision 0.91, recall 0.97 and F1 score 0.94. The former is used to label text mentions of an amino acid with a sequence number, and the latter refers to a point mutation. The remaining 18 entity types are also collected, but not processed to link them to reference terms in standardized ontologies. The codebase for the article prediction pipeline is developed in Python 3.9.12.

#### 2.1.1. Preparing the input text for prediction

Within the IUCr systems, publication text is handled as JATS (Journal Article Tag Suite; see https://jats.nlm.nih.gov/versions.html for the current version) XML files. Using XML tags such as <sec>, <sec-type> and <p>, we identified the raw text passages and section titles for each publication. The identified raw publication text was split into individual sentences. The sentences were collected and enriched with additional information such as a unique sentence identifier, a sentence character count start, which was always ‘0’, a sentence character count end, which was the total character count for a sentence, and the section a sentence belonged to. The additional information enabled tracking of sentences in the prediction pipeline and matching the predictions to specific locations within a sentence. The created sentence dictionary was used for prediction.

#### 2.1.2. Preparing the model for prediction

To efficiently run predictions with the model described in Vollmar *et al*. (2024), we applied quantisation to the original model, v2.1. During this quantisation step, we used optimum[onnxruntime] 1.2.2 to convert a 32-bit floating point model to an 8-bit integer model. Reducing the precision of the model also reduced its size from 1.3GB to 105MB. The smaller model could therefore be deployed on a CPU-only machine for inference. To assess the model’s performance after quantisation, we performed the SemEval (Segura-Bedmar *et al*., 2013) evaluation. We evaluated on an independent validation set used for benchmarking, which was published in Vollmar *et al*. (2024).

#### 2.1.3. Predicting with the model

The maximum number of tokens the model could process was 512. Therefore, we combined eight sentences into a batch and used a batch size of four (32 sentences) for prediction. For each sentence in a batch, a list of predicted named entities was returned, with each entity defined by a start and end character count. This information was used to locate the text span in the sentence, the covered text span itself, and the predicted entity type, along with a confidence score for the prediction. All predictions for a sentence were then added to the sentence dictionary created earlier under the key ‘annotations’.

#### 2.1.4. Harvesting annotations

For downstream processing of the predictions, it was necessary to modify the structure of the sentence dictionary. We chose to follow the Europe PMC annotation format as described in their instructions for annotation submission (https://europepmc.org/AnnotationsSubmission) for the final JSON file used to store the annotations. Rather than having a list of annotations for each sentence, a list of dictionaries was required with a defined set of keys:

- ‘exact’: the exact text span covered by the annotation
- ‘position’: the location of the annotation in a sentence, defined by the unique sentence identifier and the position of a word in the sentence, as given by the word count in the sentence, e.g. 25.18 refers to the 18th word in the 25th sentence
- ‘prefix’: 30 characters in a sentence preceding the exact text span
- ‘postfix’: 30 characters in a sentence following the exact text span
- ‘type”: the predicted entity type for the annotation
- ‘ai_score’: the confidence score of the algorithm for the predicted annotation
- ‘char_start’: the starting position of the exact match in a sentence
- ‘char_end’: the ending position of the exact match in a sentence
- ‘tags’: for any additional details to support the annotation, e.g. references to ontologies or controlled vocabularies; here we link to the reference sequences in UniProt

#### 2.1.5. Post-processing annotations

A number of post-processing steps were implemented to address errors in identifying text spans or assigning the correct entity type label.

A series of regular expression definitions were created to identify whether a found named entity followed expected patterns. For the entity type ‘residue_name_number’, the pattern consisted of a three-letter amino acid name and an integer for the sequence position, e.g. Arg155. If the entity type was predicted to be ‘mutant’, then the pattern was a single-letter amino acid name, sequence position, followed by another single-letter amino acid name, e.g. R155A. In cases where a single letter amino acid was found, this was expanded to the corresponding three-letter amino acid, e.g. Arg155Ala. If a found text span did not fit any of these options, e.g. due to additional characters, we tried to identify the most probable text span that would match the pattern.

To fix wrong entity type labels, we again relied on the regular expression patterns. If the pattern did not support the found entity type, we changed the label to the one that returned a regular expression match. For example, if Arg155 was labelled ‘mutant’, a search for the mutant pattern failed, whereas the one for ‘residue_name_number’ succeeded. Consequently, the entity type label was changed from ‘mutant’ to ‘residue_name_number’.

For entities of the same type that were directly adjacent based on their character counts, we combined their annotations. This step was executed as part of the prediction pipeline before writing the annotations to the Europe PMC standard JSON file.

#### 2.1.6. Enriching the sentence dictionary

Additionally, in line with the basic set of annotation requirements based on the Europe PMC standards, we added a set of optional information. These additional details covered article identifiers such as PubMed and PubMed Central identifiers, digital object identifiers (DOI), IUCr specific identifiers and the publishing license. We also added details about PDB identifiers for structures linked to a publication, the known organism with synonyms, taxonomy identifiers, and UniProt accessions for the proteins found using the PDB identifiers. All information was retrieved through API-calls either to Europe PMC using the DOI or title of an article, or to PDBe using the linked PDB identifier. The latter was possible because IUCr links and adds PDB identifiers to each of their publications, which can be easily retrieved through XML tags <ext-link>, <ext-link-type>, and <xlink:href>. ‘IUCr’ was added as a provider to keep track of who provided annotations, as they will be shared with PDBe and added to their database.

#### 2.1.7. Enriching the annotations

Following best-practices in the text mining and NLP field, the annotations were linked to references where possible. This is necessary to identify named entities without ambiguity. The two entity types of interest used here were ‘residue_name_number’ and ‘mutant’. As both entity types are protein sequence based and we know the UniProt accession for each protein found in a PDB structure, we used UniProt as our reference source. We used the SIFTS mapping files provided by PDBe which allow for a per-residue mapping between a residue in a protein structure to its equivalent residue in a UniProt accession.

Furthermore, the goal for our application was also to provide readers with experimental details to support the analysis described in the text. We therefore used the per-residue validation details provided by the PDB in the validation XML files to extract relevant statistics for each residue.

The details found for the SIFTS mapping and the validation XML were collected and attached to the ‘tags’ key of the relevant annotations, i.e. those with entity type ‘residue_name_number’ or ‘mutant’. The following details were extracted for all annotations:

- ‘pdb_id’: the PDB identifier the details belong to
- ‘pdb_res_name’: the residue name in the structure, i.e. the amino acid name
- ‘pdb_res_number’: the author provided residue number in a structure
- ‘pdb_res_seq’: the residue number in a structure as counted from the first, modelled residue
- ‘pdb_res’: a combination of the residue name and the author provided residue number in a structure
- ‘pdb_chain’: the chain identifier in a structure where the residue is found
- ‘ramachandran’: the Ramachandran score found for a residue
- ‘rotamer’: the rotamer found for a residue side chain
- ‘phi’: the phi angle found for a residue
- ‘psi’: the psi angle found for a residue
- ‘clashes’: a list of any clashes that have been found for a residue
- ‘altconf’: any alternative conformations that have been found for a residue side chain
- ‘wildtype_pdb_res’: the wildtype residue, if the entity type was “mutant”, otherwise this is empty
- ‘uniprot_id’: the UniProt accession for the protein the residue belongs to
- ‘uniprot_name’: the UniProt accession and species extension for the protein the residue belongs to
- ‘uniprot_res’: a combination of the UniProt residue name and residue number in the sequence
- ‘uniprot_uri’: the URI to the UniProt reference entry

There were additional details that are specific to the experimental method used to determine the protein structure.

For a structure determined through an X-ray diffraction experiment, we additionally provided:

- ‘rscc’: the real-space correlation coefficient found for the residue

If the structure was determined using cryo-electron microscopy, we additionally provided:

- ‘q_score’: the Q_score found for the residue

In the case of an NMR experiment used for structure determination, we additionally provided:

- ‘distances’: a list of distance outliers found for the residue
- ‘angles’: a list of angle outliers found for the residue

#### 2.1.8. Highlighting annotations in the source article

In a web environment, such as viewing a publication in a browser on a journal’s website, the ‘prefix’ and ‘postfix’ content of an annotation are sufficient in most cases to identify the sentence containing the ‘exact’ text span covered by the annotation. This text span identification follows the ‘Text Quote Selector’ approach as described in the World Wide Web Consortium (W3C) recommendation for a Web Annotation Data Model (https://www.w3.org/TR/annotation-model/).

Once annotations have been associated with an article, IUCr Journals use JavaScript (ECMAScript 2017) to retrieve the JSON data asynchronously and dynamically highlight the associated text in the HTML rendition of the article, including links to present a summary of the annotation data, as well as display the associated structure in the molecular visualization tool Mol* (Bittrich *et al*., 2024; Midlik *et al*., 2025), as described in section 2.1.9.

#### 2.1.10. Visualising residues in linked structures

Currently, IUCr Journals use the PDBe Molstar Plugin (imported from https://cdn.jsdelivr.net/npm/pdbe-molstar@3.2.0/build/pdbe-molstar-plugin.js) to display PDB structures in pop-up ‘widgets’. Highlighting a specific residue in the widget for a PDB structure is straightforward using the PDBe Molstar Plugin API. The residue can be selected using the ‘select’ method and specifying properties such as chain (e.g. struct_asym_id: ‘A’) and residue ID (*e*.*g*. residue_number: 100), along with various rendering options (*e*.*g*.representation: ‘ball-and-stick’, representationColor: {r:255,g:255,b:0}). Non-covalent interactions and volumes can be added to the focused region of the structure *via* a simple ‘interactivityFocus’ method. For further information about implementing the PDBe Molstar Plugin on web pages please see https://github.com/molstar/pdbe-molstar/wiki.

#### 2.1.10. Sharing annotations with PDBe

Annotations were designed to be shared and reused across different hardware and software platforms as we adhere to Europe PMC’s annotation JSON standard. Currently, annotation JSON files are being uploaded by IUCr Journals to a designated FTP area within the EBI infrastructure. From this location, a processing pipeline developed by the PDBe team quality checks and loads the annotations into their relational database. A set of four API endpoints (see documentation here https://www.ebi.ac.uk/pdbe/aggregated-api/ under the sections PDB and UniProt) were developed to access the annotation information in the database. The endpoints fetch database content based on a PDB ID or UniProt accession with the option to specify a particular protein chain. Furthermore, the annotations are displayed via PDBe’s newly designed webpages, which provide direct access for browsing and exploring the information when navigating through a PDB entry. Details for the webpages will be published elsewhere.

### 2.2. Hardware requirements

The annotation software will run on a modest PC *e*.*g*. 8xCore x86 CPU @ 1.00GHz + 16 GiB RAM.

### 2.3. Code availability

The code for the annotation and validation pipeline developed for this application has been published in a Github repository and is fully open access

https://github.com/PDBeurope/IUCr-annotation-pipeline.git

## 3. Results

### 3.1. Model quantisation

The model quantisation is a necessary step to improve computation time for predictions as well as to reduce hardware requirements. However, this comes at the cost of reduced prediction performance. Table 1 gives the prediction confidence of the model, either in its full or quantized version, for an unseen sentence. The quantized model displays lower values by 0.01-0.02 for the prediction confidence depending on the entity type.

**Table 1.** Confidence scores for predicted annotations in an unseen sentence for the full and the quantized model.

| start | end | text span | entity type | confidence full model | confidence quantized model |
| --- | --- | --- | --- | --- | --- |
| 1 | 23 | N-terminal acetylation | ptm | 0.937826 | 0.9305218 |
| 25 | 39 | Nt-acetylation | ptm | 0.96235585 | 0.9603966 |
| 57 | 86 | N-terminal acetyltransferases | protein_type | 0.99854547 | 0.99813604 |
| 88 | 92 | NATs | protein_type | 0.99888486 | 0.9987967 |
| 162 | 167 | Naa60 | protein | 0.9994029 | 0.9992556 |
| 180 | 184 | NatF | protein | 0.99875224 | 0.99833703 |
| 211 | 214 | NAT | protein_type | 0.99907637 | 0.99914896 |
| 229 | 253 | multicellular eukaryotes | taxonomy_domain | 0.97256976 | 0.9630969 |
| 335 | 349 | Nt-acetylation | ptm | 0.96585155 | 0.95354486 |
| 397 | 424 | lysine Nε-acetyltransferase | protein_type | 0.9983275 | 0.9971248 |
| 426 | 429 | KAT | protein_type | 0.99864644 | 0.9963988 |
| 456 | 467 | acetylation | ptm | - | 0.9865166 |
| 471 | 477 | lysine | residue_name | 0.99024945 | 0.99338126 |
| 507 | 525 | crystal structures | evidence | 0.9975256 | 0.9964277 |
| 529 | 534 | human | species | 0.99812895 | 0.9925248 |
| 535 | 540 | Naa60 | protein | 0.9994475 | 0.99933916 |
| 542 | 548 | hNaa60 | protein | 0.99933773 | 0.9990397 |
| 550 | 565 | in complex with | protein_state | 0.99842304 | 0.99808717 |
| 566 | 583 | Acetyl-Coenzyme A | chemical | 0.99915624 | 0.9989887 |
| 585 | 591 | Ac-CoA | chemical | 0.9992083 | 0.999053 |
| 596 | 606 | Coenzyme A | chemical | 0.99916434 | 0.99896336 |
| 608 | 611 | CoA | chemical | 0.999223 | 0.9989459 |
| 618 | 624 | hNaa60 | protein | 0.99931455 | 0.99901867 |
| 645 | 662 | amphipathic helix | structure_element | - | 0.9946588 |
| 645 | 656 | amphipathic | protein_state | 0.9248927 | - |
| 657 | 662 | helix | structure_element | 0.9796246 | - |
| 677 | 688 | GNAT domain | structure_element | 0.99922186 | 0.9990057 |
| 734 | 740 | hNaa60 | protein | 0.9993635 | 0.99923146 |
| 750 | 763 | β7-β8 hairpin | structure_element | 0.99896705 | 0.9989759 |
| 803 | 809 | hNaa60 | protein | 0.97456926 | 0.9877958 |
| 810 | 815 | 1-242 | residue_range | 0.89123815 | 0.949022 |
| 821 | 827 | hNaa60 | protein | 0.8546437 | 0.98467755 |
| 828 | 833 | 1-199 | residue_range | 0.85460573 | 0.91554123 |
| 835 | 853 | crystal structures | evidence | 0.9984098 | 0.9971397 |
| 899 | 905 | Phe 34 | residue_name_number | 0.99716777 | 0.99547845 |
| 1055 | 1100 | structural comparison and<br>biochemical studies | experimental_method | 0.9866565 | 0.993052 |
| 1116 | 1122 | Tyr 97 | residue_name_number | 0.9976251 | 0.99692416 |
| 1127 | 1134 | His 138 | residue_name_number | 0.9974098 | 0.9968611 |
| 1186 | 1199 | non-conserved | protein_state | 0.9988678 | 0.9984768 |
| 1200 | 1215 | β3-β4 long loop | structure_element | 0.99922025 | 0.999139 |
| 1250 | 1256 | hNaa60 | protein | 0.9993338 | 0.99913245 |

Additional assessments were carried out using the SemEval validation scheme. Tables 2-4 show the differences in performance, measured by precision, recall, and F1 score, between the full and quantized models on an unseen, independent validation set. Across this much larger benchmarking set, there is a loss of ∼0.05 for the different metrics for the quantized model compared to the full model. For the entity types ‘mutant’ and ‘residue_name_number’, which were the main focus for our pipeline, the performance changes are given in Table 5 and 6, respectively. We find that the performance of the quantized model for ‘mutant’ drops by ∼0.1 for precision, recall and F1 score. This finding comes on the back of already poor prediction performance for the full model. For ‘residue_name_number’ values, the precision, recall and F1 score are lower by 0.04-0.07, in line with the overall observed change, while the full model displays issues with overfitting, indicated by a recall of 1.

**Table 2.**
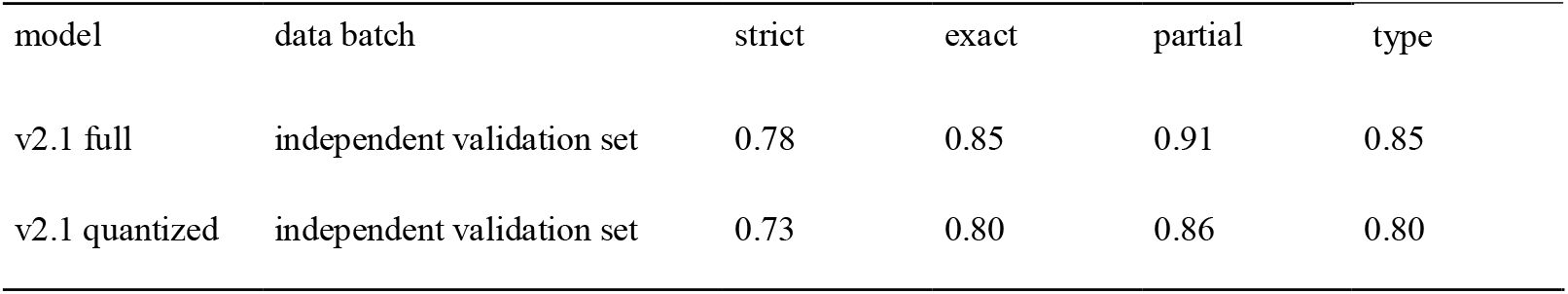
Precision determined for the full and quantized model on the respective test set and an independent validation set.

**Table 3.** Recall determined for the full and quantized model on the respective test set and an independent validation set.

| model | data batch | strict | exact | partial | type |
| --- | --- | --- | --- | --- | --- |
| v2.1 full | independent validation set | 0.75 | 0.82 | 0.87 | 0.81 |
| v2.1 quantized | independent validation set | 0.70 | 0.77 | 0.83 | 0.78 |

**Table 4.** F1 score determined for the full and quantized model on the respective test set and an independent validation set.

| model | data batch | strict | exact | partial | type |
| --- | --- | --- | --- | --- | --- |
| v2.1 full | independent validation set | 0.76 | 0.83 | 0.89 | 0.83 |
| v2.1 quantized | independent validation set | 0.71 | 0.79 | 0.85 | 0.79 |

**Table 5.** Changes of precision, recall and F1 score for the entity type ‘mutant’ for a full and quantized version of model v2.1 evaluated using SemEval and an independent validation set.

| model | strict | exact | partial | type |
| --- | --- | --- | --- | --- |
| precision full | 0.52 | 0.52 | 0.57 | 0.62 |
| recall full | 0.81 | 0.81 | 0.89 | 0.97 |
| F1 score full | 0.63 | 0.63 | 0.70 | 0.76 |
| precision quantized | 0.43 | 0.43 | 0.49 | 0.55 |
| recall quantized | 0.71 | 0.71 | 0.81 | 0.91 |
| F1 score quantized | 0.54 | 0.54 | 0.61 | 0.69 |

**Table 6.** Changes of precision, recall and F1 score for the entity type ‘residue_name_number’ for a full and quantized version of model v2.1 evaluated using SemEval and an independent validation set.

| model | strict | exact | partial | type |
| --- | --- | --- | --- | --- |
| precision full | 0.96 | 0.96 | 0.96 | 0.96 |
| recall full | 1.00 | 1.00 | 1.00 | 1.00 |
| F1 score full | 0.98 | 0.98 | 0.98 | 0.98 |
| precision quantized | 0.92 | 0.92 | 0.92 | 0.92 |
| recall quantized | 0.93 | 0.93 | 0.93 | 0.93 |
| F1 score quantized | 0.93 | 0.93 | 0.93 | 0.93 |

### 3.2. Post-processing annotations

It should be noted that although many checks have been implemented to correct errors in labelling and identifying entity types, some errors may still occur. The underlying reasons may be due to formatting errors in the JATS XML file from which the text was extracted, or the model itself may have missed a text span or assigned the wrong entity type labels. A text span may not have been encountered before and the post-processing step does not contain the necessary code to fix the error.

An example of such an error was found in ‘https://doi.org/10.1107/S205979832300311X‘. Here, the algorithm correctly identified ‘Glu89H’ (‘exact’: ‘Glu89H’) as the relevant entity in the sentence “Although Glu89H is involved in both salt bridges, the basic counter-residues are not conserved; this is because complexes A and B are nearly, but not perfectly, in twofold symmetry.”. The algorithm correctly identified the entity type, ‘type’: ‘residue_name_number’, and the prediction had very high confidence, ‘ai_score’: ‘0.9989009’. However, the post-processing step failed to remove the additional ‘H’ in this case, and the annotation does not appear on the webpages. In the preceding sentence, the named entity ‘Glu89^H_complex*B*^’ is correctly processed, removing ‘H_complexB’ and linking to its reference sequence in UniProt, as well as being displayed in Mol*.

We also found that the algorithm occasionally assigned the wrong entity type to an annotation. If an annotation was wrongly labelled ‘residue_name_number’ but was found to be of type ‘mutant’, then we changed the entity type label accordingly and set the ‘annotator’ key to ‘post_processing’ and the ‘ai_score’ key to ‘mutant’.

### 3.3. Enriching the sentence dictionary

Although the PDB has been curating protein structures for several decades, this is a semi-automated process for which requirements and standards have changed over time. At IUCr, they enriched their publications by adding links to the structures for which a publication provides a primary citation for a PDB entry. By extracting this additional information directly from the publication JATS XML, we were able to update 94 entries in the PDB with a primary citation from 35 publications, see also Predicting on IUCr journal archives.

### 3.4. Enriching annotations and mapping to references

SIFTS enabled mapping residues in a PDB entry to their counterparts in a UniProt accession. Note should be taken that the process of SIFTS-mapping only worked successfully if the authors used either the residue numbering found in the PDB structure or the numbering in the UniProt accession to refer to residues in their publication. During the mapping process, there were a few cases where the authors did not use either numbering scheme when referring to a residue in the text and therefore no link could be established. If a link was missing, we were also not able to further enrich an annotation with structure quality metrics from the validation XML file linked to a PDB entry.

### 3.5. Predicting on IUCr journal archives

All publications numbering 9152, in the journals Acta Crystallographica Section D, Acta Crystallographica Section F and IUCrJ starting from the year 2000 were submitted to the prediction and enrichment pipeline. Almost half of the publications (4124) had at least one structure linked. For the open-access publications, the predictions covered all 20 possible entity types, with those of type ‘residue_name_number’ or ‘mutant’ having undergone additional enrichment. If the publication was not open, then only predictions for ‘residue_name_number’ or ‘mutant’ were recorded. In total, 3153 publications returned a JSON file with annotations; 936 publications did not return valid annotations and 35 publications had not been recorded as a primary citation for a protein structure in the PDB, which was corrected, and annotations were added during a second round of pipeline execution. The successfully processed publications contributed a total of 781892 rows of annotations to the PDBe relational database linked to just over 10K structures.

### 3.6. Accessing the annotations in a live document

In a web environment, IUCr journals provide a number of annotations and enrichments for their publications. Which annotations will be applied can be selected by the reader. If a publication in one of the IUCr journals contains mentions for a residue with a sequence number or a point mutation, these are highlighted in the text on the webpages (Figure 1). By clicking on the highlighted residue, a selection panel will appear, which allows the selection of a structure (Figure 2). All structures that contain the highlighted residue are listed. For each listed occurrence of a residue, quality metrics are provided. Four quality metrics are provided regardless of experiment type: Ramachandran score, rotamer, phi and psi angle. For a structure determined in an X-ray diffraction experiment, we display the real-space correlation coefficient for the residue. In the case where a structure has been determined by a cryo-electron microscopy experiment, we display the Q_score and for an NMR experiment, violations for distances and angles. On clicking on the PDB entry, residue and chain label Mol* structure viewer is launched in a detached, separate window overlaying the text. It can be moved and arranged freely. This is particularly useful, if one wants to look at and compare different structures, as each PDB entry will be displayed in its own window (Figure 3).

**Figure 1.**
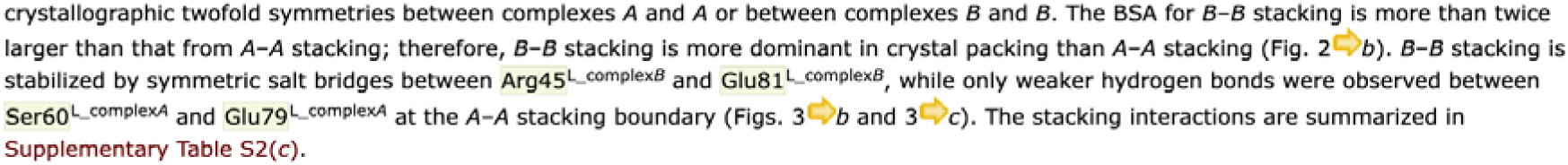
Example of highlighting residue mentions in a publication for which a link to a UniProt reference and to a residue in a structure were established.

**Figure 2.**
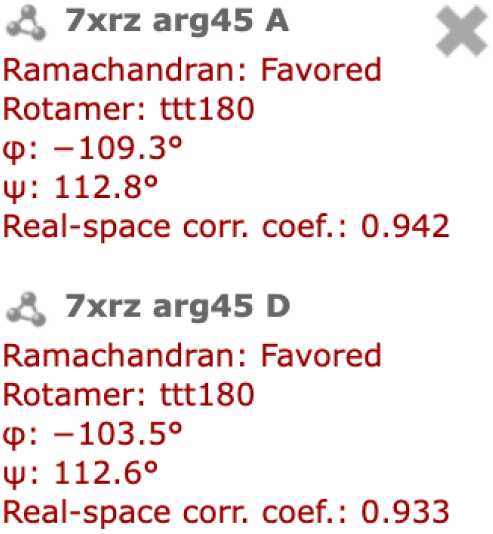
List of available structures containing the highlighted residue.

**Figure 3.**
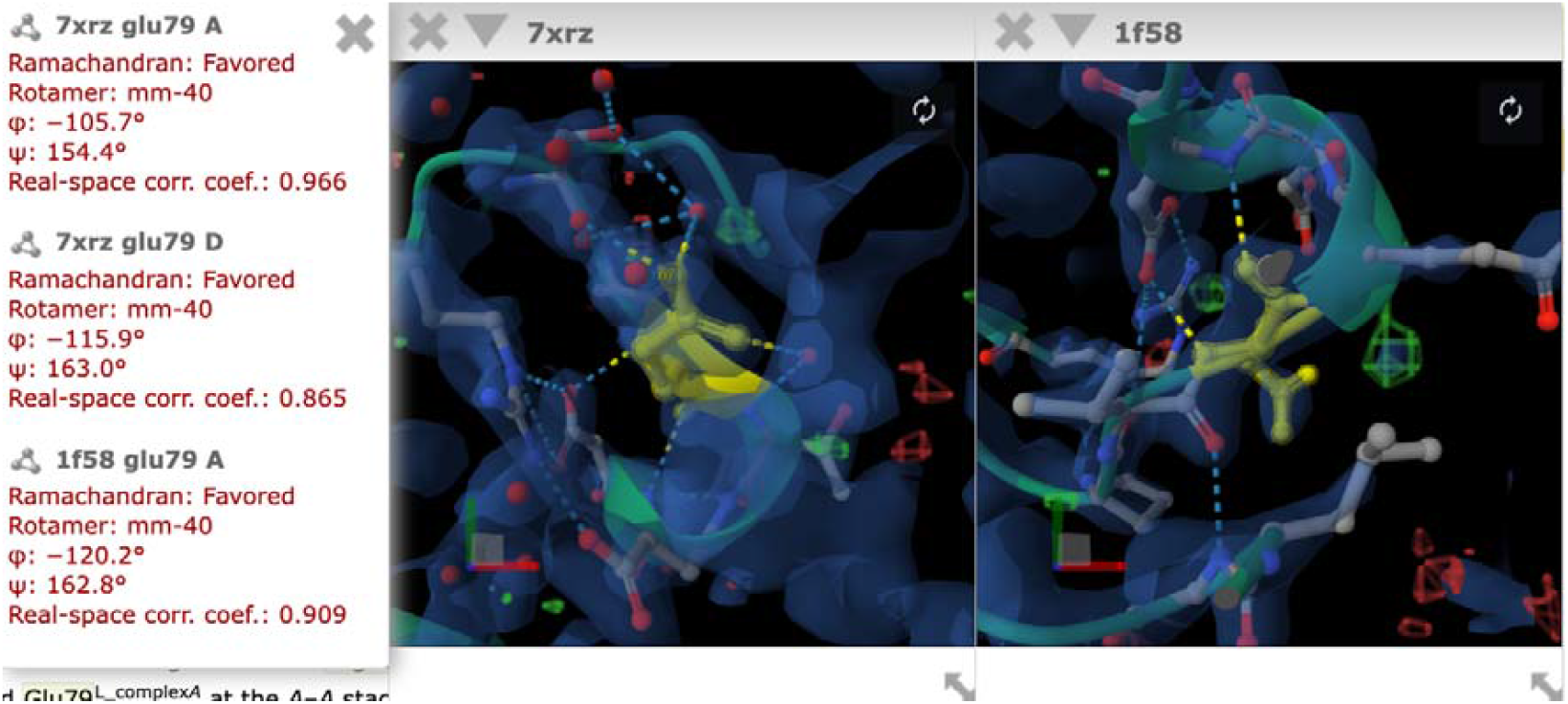
Example of how two different PDB entries can be displayed next to each other in separate Mol* windows

Residues found in the same PDB entry are displayed in the same window with the molecule centered on the selected residue. Once a molecule is displayed, it will be centered on the selected residue and chain and is displayed in cartoon style with the specific residue appearing as a ball-and-stick model coloured ‘yellow’. Found interactions, such as hydrogen bonds and salt bridges, are displayed alongside the experimental data for the selected residue and its surrounding area. For a structure derived through an X-ray diffraction experiment, an electron density map will be displayed. If cryo-electron microscopy was used to determine a protein structure, then an electric potential map is displayed. Figure 4 shows an example of how the residue and molecule are displayed. If a residue appears in several chains, like chains ‘A’ and ‘D’ in our example, then one can switch between the residues by clicking on the other one. This will not open a new window but centre the display on the selected residue instead. The window containing Mol* can be moved freely.

**Figure 4.**
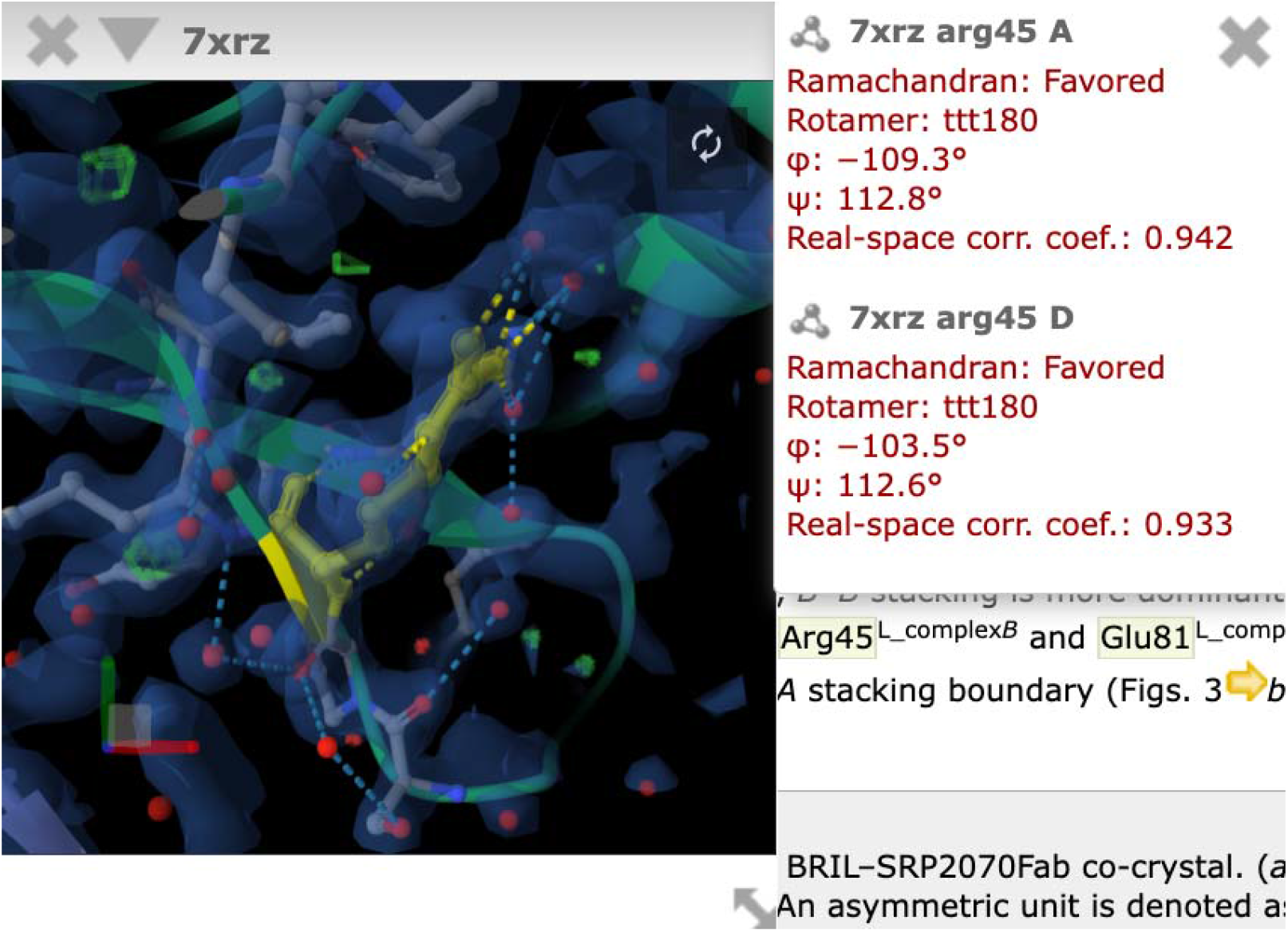
Example of how a highlighted residue is linked to a structure and displayed in Mol* molecule viewer.

## 4. Discussion

Generally, we note that the performance of the underlying model in our system is determined by the training data used during development. As was already described in Vollmar *et al*., the training data were severely imbalanced, with some entity types represented by very few examples and/or little diversity for the model to learn generalisation patterns.

To ensure the implementation of our annotation tool functions in a production environment, we quantized the model with an acceptable loss, resulting in overall performance metrics, precision, recall, and F1 score of no more than 0.05. This loss of performance allowed us to deploy a much smaller model on standard hardware. Accepting the lower performance affects the entity types ‘mutant’ and ‘residue_name_number’ differently, with the former already exhibiting low performance in the full model, which is further reduced due to quantisation. Precision measures the accuracy of positive predictions, while recall measures the model’s ability to find all relevant instances. The low precision for ‘mutant’ by the quantized model indicates that the model predicts a large number of false-positives for this entity type.

However, the recall indicates that the model can correctly identify the majority of ‘mutant’ cases in the benchmarking set.

The overall lower prediction confidence of the model resulted in a number of side effects that defined the limitations of the pipeline:

1. Entities that consist of multiple words may be predicted as a set of annotations rather than a single one
2. Entity types may have been assigned wrongly
3. Entities may have been missed by the model entirely
4. False-positive entities are predicted

The first point can be mitigated by post-processing to fuse neighbouring entities if they are of the same type, which is done as part of our prediction pipeline. The second issue we addressed through a series of regular expression checks as described in sections 2.1.5. and 3.2.

The third issue cannot be addressed through any post-processing steps. A strategy, based on the stochastic nature of these predictive models, could be to repeatedly predict on the same sentences thus increasing the chance that one of the predictions returns a result. The top-5 predictions can then be used to determine the most likely result based on a majority count for confidence and entity type. Alternatively, a user feedback mechanism could be developed through IUCr and PDBe pages to identify missing annotations. The latter would also be beneficial to identify other errors found in annotations. Aside from the model not finding a named entity in the text, an annotation can also be missing because the downstream pipeline was not able to link to a sequence in the structure or a UniProt accession. This is usually the case if the authors’ used a sequence numbering to refer to a residue in the text that does not reflect either of the resources, PDB or UniProt. In such cases, no linking was possible and the annotation will be missing from the IUCr pages, the PDBe relational database, API endpoints and the PDBe webpages.

The last issue, having a high false-positive rate as we find for ‘mutant’, can in some cases be traced back to having been assigned the wrong entity type and overlaps with issue three.

Additionally, when developing the predictive model, the definition of ‘mutant’ used was not limited to point mutations, as described in Section 2.1.5. but covered any form of sequence alteration, like removing anything from a few residues up to whole domains. For example, in the sentence “The accumulation of SRP2070Fab molecules by stacking is remarkable and is consistent with the finding that stacking of SRP2070Fab is predominant in known crystal structures of BRIL-fused GPCRs complexed with SRP2070Fab.” from ‘https://doi.org/10.1107/S205979832300311X‘above, ‘BRIL’ is here not identified as ‘protein’ entity type but as ‘mutant’. This is correct, as in this context, BRIL, although being a protein, is also artificially fused to a GPCR and is therefore also of the ‘mutant’ type in the wider sense of our definition. Crucially, we not only check whether an identified entity follows a regular expression pattern for ‘mutant’ or ‘residue_name_number’ but we also check whether the wordspan for an entity type can be matched to a residue via SIFTS downstream. If the textspan cannot be resolved to match a residue, then the annotation is discarded. In the sentence “The gene encoding a Tobacco etch virus (TEV) protease cleavage site and BRIL (Ala1–Leu106) was synthesized and subcloned into pET-28a(+) using BamHI and HindIII.” ‘pET’ is predicted to be of entity type ‘mutant’. We can see that this text span doesn’t match what we would expect for a point mutation or even a deletion, but some of the context in the sentence points towards sequence alterations more generally. However, this prediction comes with a confidence level of ‘ai_score’ 0.46529654, indicating that the model is not very confident in the accuracy of the prediction. Setting a prediction confidence threshold would therefore be another option to filter annotations and exclude false-positives.

We successfully processed 76% of the publications in the entire IUCr archive for three journals, dating back to 2000. The remaining 24% could not be processed for several reasons:

1. Publications may have been linked to a structure, but there was no mention of specific residues in the text
2. The authors of the publication may not have used a numbering scheme for residues mentioned in the text that could be linked to either the PDB structure or a UniProt accession.
3. In some cases, our pipeline failed entirely.

Issues one and two are inherent to the specific publication and cannot be fixed through any checks and quality control. Therefore, no annotations will be generated, and although only spot testing was done for failure cases, these represent the majority of failed publications. For the third issue, we will investigate the reason for failure for the particular publication and, if possible, improve our pipeline. There may be text spans that we are currently unable to process and analyse. But also, during several steps, we execute API calls to other resources. If these resources are offline for reasons out of our control, then our pipeline will create its usual output but without any content.

## 5. Conclusion

Here, we present a novel system that enables the linking of text mentions of functionally important residues to their corresponding counterparts in a linked protein structure and the associated UniProt accession. Within the web-based publication environment at IUCr these links are used to highlight functionally important residues in their text context, while a molecular viewer can be opened to view the respective residue in its structural context, supported by experimental quality metrics. We extracted the annotations and shared them with PDBe for indexing and open sourcing, allowing for downstream analysis, further linking on their webpages, and dissemination to the wider life science community.

## Acknowledgements

We thank Adam Midlik from the PDBe team for his support in implementing the Mol* viewer for use by IUCr Journals. Funding was provided to M.V. as an ARISE Fellowship from the European Union’s Horizon 2020 research and innovation programme under the Marie Skłodowska-Curie grant agreement No. 945405.

